# Single cell chemotactic responses of *Helicobacter pylori* to urea in a microfluidic chip

**DOI:** 10.1101/045328

**Authors:** Xuan Weng, Suresh Neethirajan, Adam Vogt

**Author notes:** (X. W.). (A.V.).

## Abstract

*Helicobacter pylori* is a spiral-shaped bacterium that grows in the human digestive tract; it infects ~50% of the global population. *H. pylori* induce inflammation, gastroenteritis, and ulcers, which is associated with significant morbidity and may be linked to stomach cancer in certain individuals. Motility is an essential virulence factor for *H. pylori*, allowing it to migrate toward and invade the epithelial lining of the stomach to shelter it from the harsh environment of the stomach. *H. pylori* senses pH gradients and use polar flagella to move towards the epithelium where the pH approaches neutrality. However, its chemotaxis behaviors are incompletely understood. Previous in vitro tests examining the response of *H. pylori* to chemical gradients have been subjected to substantial limitations. To more accurately mimic/modulate the cellular microenvironment, a nanoporous microfluidic device was used to monitor the real time chemotactic activity of single cell of *H. pylori* in response to urea. The results showed that microfluidic method is a promising alternative for precisely studying chemotactic behavior of bacteria.

## 1. Introduction

*Helicobacter pylori (H. pylori)* is a common bacterium that was first identified in 1983 [1]. *H. pylori* cells are adapted to live in the harsh, acidic environment of the stomach and their spiral shape allows them to penetrate the stomach lining. *H. pylori* colonizes the stomachs of approximately 50% of the human population and contributes to overt gastric disease, peptic inflammation, and ulceration. *H. pylori* also have an etiological relationship with gastric cancer [2]; there is a gut microbe-host relationship that exists between *H. pylori* and the human or animal gut. The ability of *H. pylori* to interact with cells within the gastric epithelium is greatly influenced by its motility [3,4]. Therefore, investigating chemotaxis in *H. pylori* has significant applications in food safety, the human health sector, and the farming/agricultural industry.

In practice, first-line therapies fail 10-20% of the time [5, 6], in part due to antibiotic resistance and poor compliance with complicated dosing schedules [7]. Motility in *H. plyori* is essential for initiating and maintaining infections in humans and animal models [8]. Therefore, motility is a major virulence factor, and studies have reported that non-motile *H. pylori* cells are less virulent [9]. Therefore, chemicals that modulate motility play an important role in virulence and are able to affect the severity of symptoms during *H. pylori* infection [9]. *H. pylori* cells utilize multiple flagella to move through the gastric environment, a process which is governed by chemotaxis (typically towards areas of lower pH) [10,11].

Chemotaxis is defined as the preferential movement towards more favorable chemical environments. In the growth cycle of *H. pylori* within the gastric mucosal layer, chemotaxis-directed motility is a remarkable characteristic [9]. Chemotaxis helps *H. pylori* to establish and maintain infection in animal models [10]. Although a significant amount of work has been done to measure chemotaxis and general motility in *H. pylori*, there is no consensus on how these processes work during infection. For example, Abdollahi and Tadjrobehkar [9] and Worku et al. [12] used very similar experimental methods, but obtained opposite findings in regards to the response of *H. pylori* to aspartate. Using in vivo models,Williams et al. [13] demonstrated that *H. pylori* uses chemotaxis to guide itself to the stomach epithelium. Many other studies have assessed the response of *H. pylori* to gradients of physiologically relevant compounds in vitro [14,15]. However, the experimental conditions in those studies do not accurately mimic the conditions present in vivo. In particular, these models do not incorporate a viscoelastic medium or pH gradient, which are important environmental conditions relevant to infection [16].

It has been demonstrated that the pathogenic mechanisms of *H. pylori* are influenced by urea. Urease activity allows *H. pylori* cells to survive in an acidic gastric environment by creating a "cloud" of ammonia around the bacterium that helps to neutralize the pH of the surrounding environment, thus playing an important role in virulence and pathogenesis of gastric and peptic ulcers; ammonia generation leads to the destruction of the gastric epithelium [17, 18]. Furthermore, studies show that cytoplasmic urease plays a critical role in the chemotactic behaviors of *H. pylori* [19, 20].

Traditionally, microbes have been studied in the laboratory as cultures of isolated species using conventional microbiological techniques, in part due to the complexities involved in designing and building physiologically relevant environments. Microfluidics overcomes the limitations of traditional methods, and offers techniques for cultivating and investigating microbes in a more clinically realistic and physiologically relevant environment. Microfluidic devices have been become powerful tools used in cell biology applications, including cell culture [21] and handling and analysis [22, 23]. These devices allow researchers to better mimic in vivo physiological conditions due to their unique properties of the microenvironment control, customizable microstructure, and perfusion controllability [24]. Microfluidics provides a large surface area to volume ratio compared to standard cell culture plates due to their micrometer dimensions. The small dimensions of the microchannels facilitate the formation of laminar flow, such that the liquid–liquid interface mass transport is realized by diffusion. This feature also establishes microfluidic systems as tools for mimicking the in vivo environment.

In addition, single cell *H. pylori* chemotaxis has not been studied mainly due to the challenges associated in culturing the cells under anaerobic conditions. Although preferring microaerophilic conditions, *H. pylori* has the ability to grow equally well in vitro under microaerobic or aerobic conditions, if cultured at high bacterial concentrations [25]. With the advent of microfluidic technology and with instrumentation, it is possible to create a conducive microenvironment to study the chemotaxis of cells in their natural state. Microfluidics is reportedly much more sensitive when measuring chemotaxis in *Escherichia coli* than the capillary and soft agar assays traditionally used with *H. pylori* [26]. The same order of the size of the cells and microchannel enables to observe the cells and the subtle movements of single cells at high resolution. Currently there are no reports where microfluidics has been used to study *H. pylori.* In this study, the single cell swimming dynamics of *H. pylori* were investigated using a combination of imaging and microfluidic systems. Results generated with our methodology aids in improving the understanding of the behavior of this pathogenic bacterium and also provide a promising way to investigate urease inhibition in regards to the treatment of *H. pylori* infection.

## 2. Materials and Methods

### 2.1 Bacterial strains and cell culturing

*H. pylori* strain SS1 (provided as a gift from The Hospital for Sick Children, Toronto, Canada) was streaked on 5% horse blood agar plates (Oxoid/ Thermo fischer number MP2300). Plates were then incubated in a jar filled with premixed gas (5% O2, 10% CO2, balance N2) at 37°C for four days. A single colony was inoculated into a test tube with Brucella broth and incubated in the premixed gas filled jar at 37°C for one to four days. Before an assay, the bacteria culture was washed with chemotaxis buffer and pelleted by centrifugation (SciLogex D3024, Berlin, CT) at 3000 rpm for 3 min, a total of twice. The pellet was then gently resuspended in chemotaxis buffer for further use. Potassium phosphate buffer of 10 mM (pH 7.0) served as the chemotaxis buffer and washing solution.

All of the chemicals (urea, sodium and potassium bicarbonate) and Brucella culture medium used for bacterial growth were purchased from Sigma Aldrich Canada.

### 2.2. Microfluidic platform and device fabrication

The details for the design of the microfluidic device used in this study have been described in our previous study [27]. Briefly, the microfluidic chip consists of one central (chemotaxis) channel (415 μm in width) with a viewing area (200 μm in width) in the middle and two side channels (280 μm in width) for feeding solution. The side channels have narrow neck of 40 μm in the middle. The “neck” is connected with the viewing area by hundreds of nanoporous structure membranes (800 nm in diameter) that are used for solution diffusion. The depth of the microfluidic chip is 10 μm.

The silicon and polymer master was fabricated using a combination of electron beam lithography, anisotropic silicon etching, and cross linking of the polymer [26]. The PDMS microfluidic chip was made by using a master mold and standard soft lithography. Briefly, a degassed mixture of PDMS prepolymer and curing agent (10:1 w/w, Sylgard, Dow Corning, Burlington, ON, Canada) was poured over the top of the silicon mold and baked for 4 hours at 75°C to harden the PDMS. The PDMS replica was then peeled off the master mold, punched to form the inlets and outlets, and bonded onto a glass slide (25 × 75 × 1 mm, VWR International, Suwanee, GA, USA) after plasma cleaning (Harrick Plasma, Ithaca NY) for 40 s. Finally, the bonded chip was baked for 30 minutes at 60 °C to solidify the bond.

The gradient generation test of the microfluidic device was also carried out as described previously[27]. Two fluorescent dyes (5 μm Fluorescein and 5 μm Texas Red) were used in the test to determine the diffusion rate. A rapid concentration gradient was clearly observed using the liquid–liquid interface.

### 2.3. Experimental setup

In each assay, a blocking step was performed to avoid the nonspecific binding of bacteria to the surfaces by filling the microchannel with 0.1% bovine serum albumin (BSA) for 10 min and washing with distilled water.

To begin, we determined the baseline (control) motility of *H. pylori.* After blocking, bacterial solution was manually dispensed into the central channel with a 1 ml syringe. Chemotaxis buffer was dispensed into the side channels using a syringe pump (Chemyx Fusion Touch, Stafford, TX) to maintain a constant flow rate of 30 μl per hour. After the viewing area filled with bacterial solution, the chip stood for a couple of minutes to stop mechanical flow so that only bacterial self-propelled motion would be present in the viewing chamber. The inlet and outlet of the central channel was then plugged with PDMS stoppers. A video of the viewing area was recorded. Similarly, for the chemotaxis assay, after filling the chamber with bacterial solution, chemotaxis buffer and urea solution were dispensed into the side channels using a syringe pump (Chemyx Fusion Touch, Stafford, TX) to maintain a constant flow rate of 30 μl per hour. A video of the viewing chamber was recorded.

### 2.4. Imaging processing and analysis

Videos of the viewing chamber were taken with a Nikon Eclipse Ti inverted microscope mounted with a Nikon DS-QiMc microscope camera and a Nikon NIS Elements BR version 4.13 software (Nikon Instruments Inc., Melville, NY). The video was captured at 640 × 512 resolution with a 10 ms exposure, 2× analog gain, and 15 fps in 30 s-long sections.

Image processing and data analysis was carried out using the public domain program ImageJ (http://rsb.info.nih.gov/ij/). The taken video was first divided into 10 s clips that were processed individually. Twenty-thirty bacteria were selected for tracking in each test. Tracking was performed on single bacteria, frame by frame, using the Manual Tracking Plugin (Fabrice Cordelires, Institut Curie, Orsay France). Data obtained were then exported to the Chemotaxis and Migration Tool (Ibidi Software, Munich, Germany) for further analysis. The cell movement was characterized by microscopic imaging, cellular velocity analysis, and directness analysis, which refers to the linearity of cell trajectories calculated by comparing the length of the line segment connecting the starting and ending points of the cell's trajectory to the total length travelled by the cell.

## 3. Results and discussion

Chemotaxis aids *H. pylori* in nutrient acquisition, allowing it to persist within the mammalian host [15]. Urea is synthesized in the liver, circulated in the bloodstream, and diffuses into the gastric lumen via capillaries, which generates concentration gradients of urea across the mucous layer [19]. In this study, we conducted two types of chemotaxis assays: spatial assays observing bacterial migration to a particular area in response to a urea gradient and temporal assays monitoring the flagellar switching of a single cell. The most popular assay to investigate the migration of *H. pylori* is the soft-agar method [28], which requires rich media and serum to support growth and does not evaluate the effect of a single chemotactic agent; this method assesses only general chemotactic ability [15]. Using our microfluidic device, a concentration gradient of urea was generated in the microchannel [27], allowing us to examine the response of *H. pylori* to a single chemoeffector. Another advantage of our microfluidic device is that the small scale of the observation area allows for the surveillance of the subtle movements of a single cell at high resolution. We thus investigated the chemotactic activities of a single bacterium, including its complex bacterial swimming behavior and the frequency and type of directional changes or tumbles.

### 3.1. Impact of urea on H. pylori cellular velocity

Twenty cells were randomly selected and tracked for at least 10 s to analyze individual bacterial swimming behavior. Cellular velocity is shown in Figure 1. At baseline (no urea added), the cells exhibited a low velocity with a large standard derivation. The relative limited motility of *H. pylori* in chemotaxis buffer may be attributed to the lack of an adequate internal energy store to power the flagella [28]; chemotaxis buffer is nutrient poor. The relatively large standard deviation for cellular velocity reflects the instability in *H. pylori* motility, which is partially the result of a frameshift mutation-prone repetitive sequence in fliP, the flagellar biosynthetic gene [29]. A decrease in the cellular velocity was observed (compared to baseline) when concentration gradients of urea were generated in the microenvironment. However, there were no significant differences in velocity between cells exposed to 10 versus 100 mM urea.

**Figure 1.**
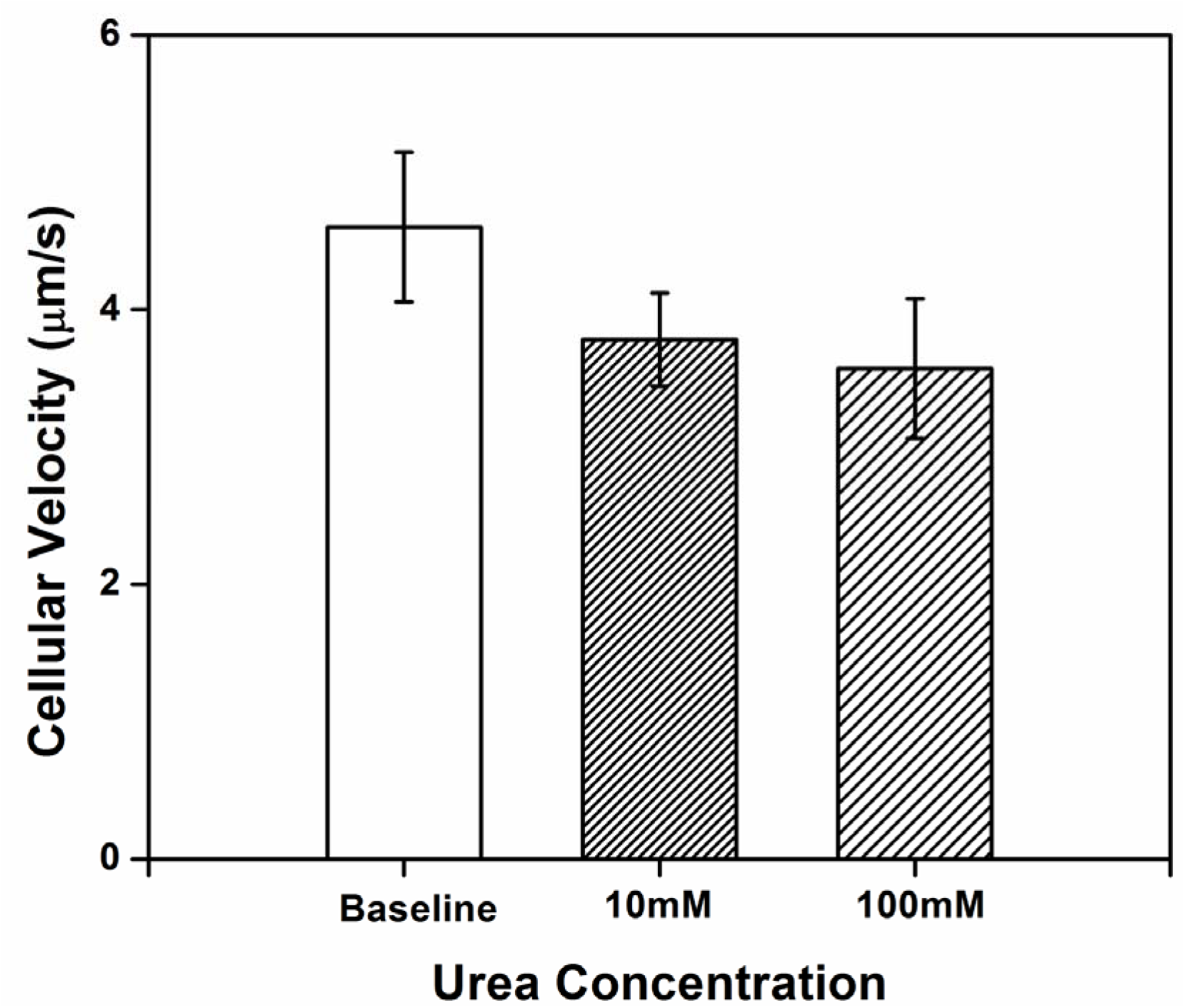
Cellular velocity of *H. pylori* in chemotaxis buffer with 10 and 100 mM urea within the microfluidic device. An obvious drop was observed between baseline and 10 or 100 mM urea-exposed cells. Three independent tests were conducted and 20-30 bacteria were randomly selected for tracking and analyzing in each test.

### 3.2. Impact of urea on H. pylori directness

Directness was used as a measure of how often the cell changed its travelling path and the randomness of movement, comparing the actual distance covered by the cell to the straight line distance between the starting point and the end point of the cell's movement. The high directness value obtained reflects the inherently random movement of the *H. pylori* cells in chemotaxis buffer. Figure 2 shows the directness of *H. pylori* movement when exposed to concentration of 10 and 100 mM urea within the microenvironment. Compared to baseline, the randomness of *H. pylori* cellular movement decreases with exposure to urea. The twirling and tumbling behaviors exhibited in chemotaxis buffer without urea are practically absent when urea is present. Sparse tumbling was observed with urea, but this was only in the direction of the urea source.

**Figure 2.**
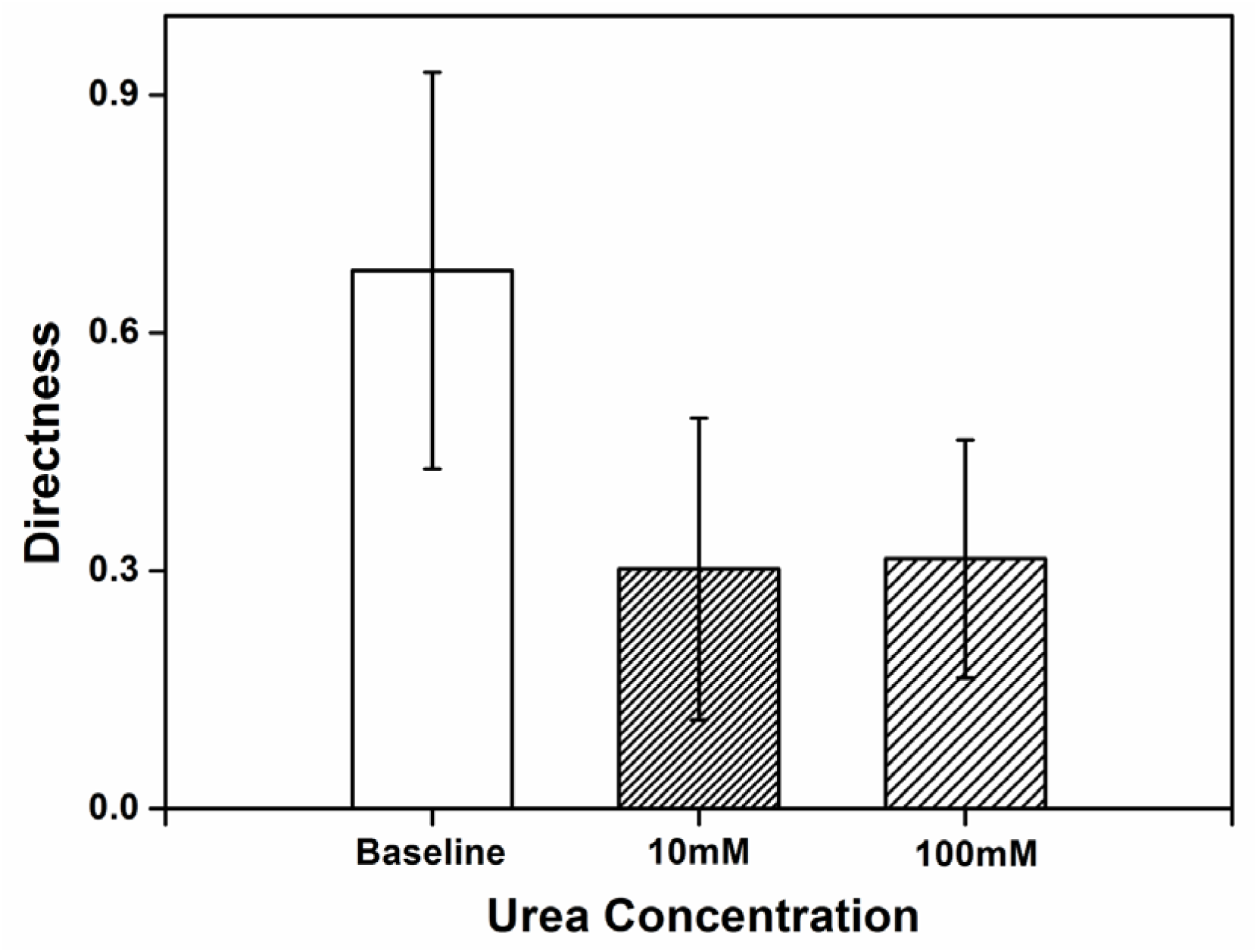
Directness of the *H. pylori* in chemotaxis buffer with 10 and 100 mM of urea within the microfluidic device. A steep decrease was observed in the directness when urea solution was introduced, while no significant differences were observed between 10 and 100 mM urea-exposed cells. Three independent tests were conducted and 20-30 bacteria were randomly selected for tracking and analyzing in each test.

### 3.3. Impact of urea on the type and frequency of H. pylori movements

The cell dynamics and mechanics of *H. pylori* during motility in chemotaxis buffer were also characterized by analyzing the video recorded by the microscope at slower acquisition recording rates. Most bacterial cells exhibited twirling and tumbling behavior in chemotaxis buffer. Figure 3 gives an example of the typical motility behavior of a single cell. Three dimensional rotations were observed as indicated by arrows shown in the images.

**Figure 3.**
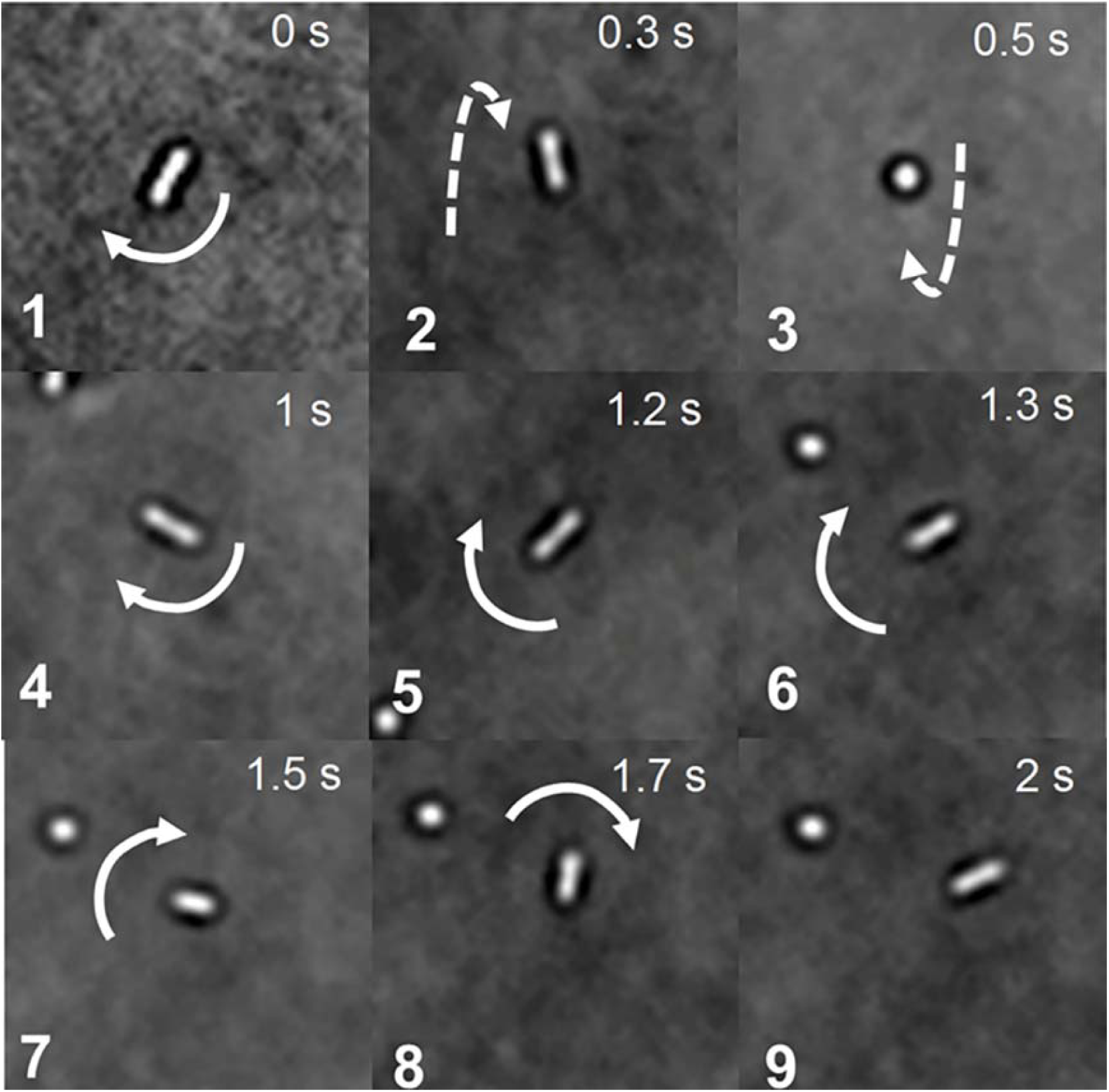
Images of *H. pylori* in chemotaxis buffer within a time frame of 2 s, which shows that the cells exhibit complex moving behavior, including twirling and tumbling. The motility was documented using a phase contrast video microscope with a 60× lens at room temperature. Arrows indicate the direction of the movement of the cell. Frame 2 and 3 show a tumbling in z direction. Frames 6-9 show a tumbling in x-y direction.

The chemotactic behavior of *H. pylori*, dictated by flagellar locomotion, is a crucial factor in the bacterial colonization process [19]. *H. pylori* cells normally have 2-6 sheathed flagella that the bacterium uses to move to and within its ecological niche. The *H. pylori* flagella consist of three structural components: a basal body, an external helically shaped filament, and a hook. These components contain the proteins required for flagellar rotation and chemotaxis, which function as a propeller when rotated at its base [30]. Nakamura et al. [19] have suggested that *H. pylori* utilize a proton motive force for flagellar motility. It has also been reported that the velocity of *H. pylori* cells decreases in response to increasing concentrations of urea, likely due to lack of protons that would be required to neutralize excess ammonia when urea is present in a more neutral (pH 4.5-7) environment [31]. In vitro tests show that *H. pylori* is acid-sensitive, but is protected from acid (pH < 4.0) by the ammonia released during urea hydrolysis, which is associated with its inherent urease activity [32]. Therefore, *H. pylori* can survive in the presence of urea when exposed to acid in vitro. In keeping with these observations, we found that *H. pylori* exhibits restricted motility in a near neutral environment.

*H. pylori* is generally thought to have four chemoreceptors: TlpA, TlpB, TlpC, and TlpD. Proteins TlpA, TlpB, and TlpC span the inner membrane, which means that part of the protein is located within the periplasm, which is the space between the two membranes of Gram-negative organisms, while another part of the protein is located in the cytoplasm; TlpD is localized completely within the cytoplasm [15]. In *H. pylori*, four different regulatory proteins, CheA, CheY, CheW, and CheZ, are responsible for the transduction of the signal that initiates flagellar motor. When compounds, such as urea bind to the Tlp receptor, allosteric changes in the receptor propagate to a CheY kinase, called CheA. CheA then catalyzes the transfer of a phosphate from adenosine triphosphate (ATP; cellular energy source) to a cytoplasmic protein, CheY; the phosphorylated form of CheY is called CheY-P. CheY-P is able to interact with the motor at the base of the flagellum. The flagellar motor essentially operates as a sensor for CheY-P, whose output is a tendency toward rotation of the flagellum leading to cell tumbling [30]. The interaction of physically proximal receptors may be an important factor in deciding how cells react to even small changes in chemoattractant concentrations.

In our study, *H. pylori* cells were attracted to urea. Figure 4 gives an example how the cells moved towards the side of the microfluidic device where the concentration of urea was greatest. The chemotactic activities exhibited by *H. pylori* presented in response to urea suggest that the proton motive force generated in the process is adequate for swimming in chemotaxis buffer. As discussed, adaptive mechanisms are responsible for the chemotactic movement toward attractants [30, 33]. In *H. pylori*, chemoattractant sensing is mediated by methyl-accepting chemotaxis proteins (MCP), which contain a large cytoplasmic signaling and adaptation domain. When an attractant binds to an MCP, the activity of CheA is inhibited, resulting in increased methylation of the Tlp receptor by CheR. Consequently, adaptation of the receptor returns signaling to a state similar to that prior to stimulus challenge, which generates periodical “running” induced by dephosphorylated CheA and CheY [30]. Attractant-bound MCP may suppress CheA activity, reducing the pools of CheY-P, which may explain why the *H. pylori* exhibited less frequent tumbling when exposed higher concentrations of urea. Ultimately, this may help to explain how *H. pylori* senses concentration gradients of urea to move towards higher concentrations. In combination with the restricted motility discussed earlier, this helps to explain the lower directness (reflecting less random movement).

**Figure 4.**
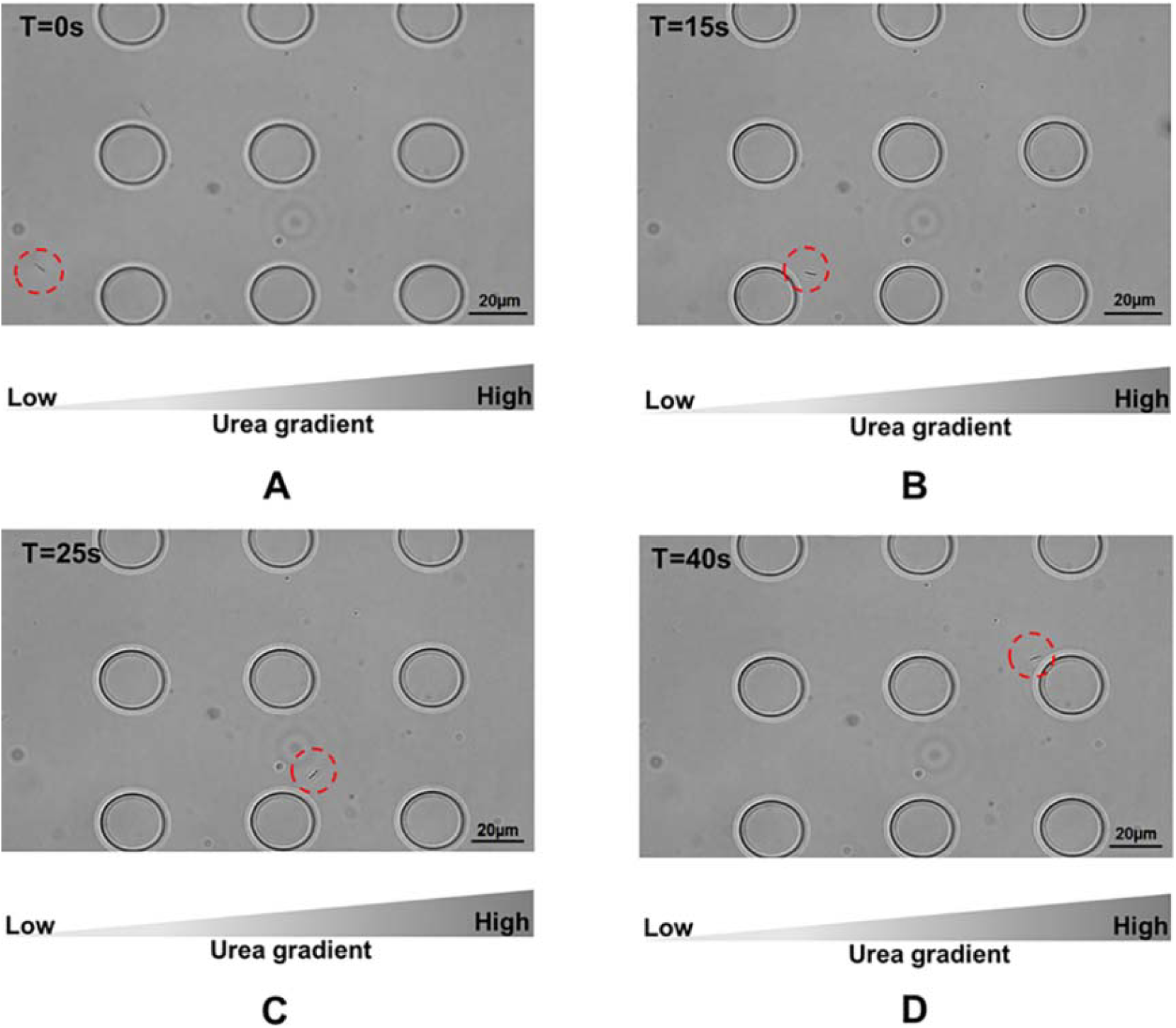
Time-lapse migration image of an *H. pylori* cell response to a gradient concentration of urea solution within the microfluidic device (a low to high concentration gradient of urea was generated within the microfluidic device along the left hand side to the right). Red circles labeled in the picture indicates the tracked cell's location at the time points of (A) 0 s, (B) 15 s, (C) 25 s, and (D) 40 s, respectively.

## 4. Conclusions

In conclusion, chemotaxis toward urea was observed when *H. pylori* was monitored in the near neutral micro-environment established within our nanoporous microfluidic chip designed for observing bacterial movement. *H. pylori* exhibits chemotactic behaviors in response to specific stimuli (e.g. urea), which is governed by the proton motive force. These behaviors were characterized by a decrease in both the velocity and random motion of the cells. Here, the urease-driven hydrolysis of intracellular urea may contribute to the generation of the proton motive force, which powers rotation of the *H. pylori* flagellum motor. Classically, *H. pylori* motility behaviors have been difficult to study by conventional methods. Therefore, the use of a microfluidic chip to mimic the ecological niche of this bacterium is a promising alternative for studying its chemotactic behavior, as evidenced by our findings.

## Acknowledgements

This study is supported by grants from the Natural Sciences and Engineering Research Council of Canada (NSERC). The authors sincerely thank Dr. Nicola Jones of the Hospital for Sick Children for providing the *Helicobacter Pylori* bacterial strains for our study.

## Author Contributions

In this study, Suresh Neethirajan conceived and designed the experiments. Xuan Weng and Adam Vogt conducted the described research and performed the processing and analysis of the experimental results.Xuan Weng prepared the manuscript. Suresh Neethirajan revised and proofread the manuscript.

## Conflict of Interest

The authors declare no conflict of interest.

## References

1. Warren, J. R.; Marshall, B. Unidentified curved bacilli on gastric epithelium in active chronic gastritis. Lancet 1983, 321, 1273–1275.

2. Holmes, E.; Li, J. V.; Athanasiou, T.; Ashrafian, H.; Nicholson, J. K. Understanding the role of gut microbiome-host metabolic signal disruption in health and disease. Trends Microbiol. 2011, 19, 349–359.

3. Keilberg, D.; Ottemann, K. M. How Helicobacter pylori senses, targets, and interacts with the gastric epithelium. Environ. Microbiol. 2016, DOI: 10.1111/1462-2920.13222. Available online: 4 February 2016.

4. Aihara, E.; Closson, C.; Matthis, A.L.; Schumacher, M.A.; Engevik, A.C.; Zavros, Y.; Ottemann, K.M., Montrose, M.H. 2014. Motility and chemotaxis mediate the preferential colonization of gastric injury sites by Helicobacter pylori. PLoS Pathog, 2014, 10, e1004275.

5. Fischbach, L. A.; Goodman, K. J.; Feldman, M.; Aragaki, C. Sources of variation of Helicobacter pylori treatment success in adults worldwide: a meta-analysis. Int. J. epidemiol. 2002, 31, 128–139.

6. Chuah, S.; Tsay, F.; Hsu, P.; Wu, D. A new look at anti-Helicobacter pylori therapy. World J. Gastroenterol. 2011, 17, 3971–3975.

7. Graham, D. Y. Helicobacter pylori eradication therapy research: Ethical issues and description of results. Clin. Gastroenterol. Hepatol. 2010, 8, 1032–1036.

8. Rust, M.; Schweinitzer, T.; Josenhans, C. Helicobacter flagella, motility and chemotaxis. In Helicobacter pylori: Molecular Genetics and Cellular Biology, Yamaoka, Y., Eds; Horizon Scientific Press, UK, 2008, Chapter 4, pp. 61–86.

9. Abdollahi, H.; Tadjrobehkar, O. The role of different sugars, amino acids and few other substances in chemotaxis directed motility of helicobacter pylori. Ira. J. basic med. Sci. 2012, 15, 787–794.

10. Terry, K.; Williams, S. M.; Connolly, L.; Ottemann, K. M. Chemotaxis plays multiple roles during Helicobacter pylori animal infection. Infect. Immune. 2005, 73, 803–811.

11. Foynes, S.; Dorrell, N.; Ward, S. J.; Stabler, R. A.; McColm, A. A.; Rycroft, A. N.; Wren, B. W. Helicobacter pylori possesses two CheY response regulators and a histidine kinase sensor, CheA, which are essential for chemotaxis and colonization of the gastric mucosa. Infect. Immun. 2000, 68, 2016–2023.

12. Worku, M. L.; Karim, Q. N.; Spencer, J.; Sidebotham, R. L. Chemotactic response of Helicobacter pylori to human plasma and bile. J. Med. Microbiol. 2004, 53, 807–811.

13. Williams, S. M.; Chen, Y. T.; Andermann, T. M.; Carter, J. E.; McGee, D. J.; Ottemann, K. M. Helicobacter pylori chemotaxis modulates inflammation and bacterium-gastric epithelium interactions in infected mice. Infect. Immune. 2007, 75, 3747–3757.

14. Lertsethtakarn, P.; Ottemann, K. M.; Hendrixson, D. R. Motility and chemotaxis in Campylobacter and Helicobacter. Annu. Rev. Microbiol. 2011, 65, 389–410.

15. Lertsethtakarn, P.; Draper, J.; Ottemann, K. M. Chemotactic Signal Transduction in Helicobacter pylori. In Tivo-component Systems in Bacteria, Gross, R., Beier, D., Eds.; Horizon Scientific Press, UK, 2012, Chapter 17, pp. 355.

16. Sycuro, L. K.; Wyckoff, T. J.; Biboy, J.; Born, P.; Pincus, Z.; Vollmer, W.; Salama, N. R. Multiple peptidoglycan modification networks modulate Helicobacter pylori’s cell shape, motility, and colonization potential. PLoS Pathog. 2012, 8, e1002603.

17. Dunn, B. E. Pathogenic mechanisms of Helicobacter pylori. Gastroenterol. Clin. North Am. 1993, 22, 43–57.

18. Follmer, C. Ureases as a target for the treatment of gastric and urinary infections. J. Clin. Pathol. 2010, 63, 424–430.

19. Nakamura, H.; Yoshiyama, H; Takeuchi, H.; Mizote, T.; Okita, K.; Nakazawa, T. Urease plays an important role in the chemotactic motility of Helicobacter pylori in a viscous environment. Infect. Immun. 1998, 66, 4832–4837.

20. Umamaheswari, R. B.; Jain, S.; Tripathi, P. K.; Agrawal, G. P.; Jain, N. K. Floating-bioadhesive microspheres containing acetohydroxamic acid for clearance of Helicobacter pylori. Drug Deliv. 2002, 9, 223–231.

21. Halldorsson, S.; Lucumi, E.; Gömez-Sjöberg, R.; Fleming, R. M. Advantages and challenges of microfluidic cell culture in polydimethylsiloxane devices. Biosens. Bioelectron. 2015, 63, 218–231.

22. DiCicco, M.; Neethirajan, S. An in vitro microfluidic gradient generator platform for antimicrobial testing. BioChip J. 2014, 8, 282–288.

23. Wright, E.; Neethirajan, S.; Weng, X. Microfluidic wound model for studying the behaviours of Pseudomonas aeruginosa in polymicrobial biofilms. Biotechnol. Bioeng. 2015, 112, 2351–2359.

24. Terry, J.; Neethirajan, S. A novel microfluidic wound model for testing antimicrobial agents against Staphylococcus pseudintermedius biofilms. J. Nanobiotechnology 2014, 12, 1.

25. Bury-Moné, S.; Kaakoush, N. O.; Asencio, C.; Mégraud, F.; Thibonnier, M.; De Reuse, H.; Mendz, G. L. Is Helicobacter pylori a true microaerophile? Helicobacter 2006, 11, 296–303.

26. Englert, D. L.; Jayaraman, A.; Manson, M. D. Microfluidic techniques for the analysis of bacterial chemotaxis. In Chemotaxis. Humana press. 2009. pp. 1–23.

27. Wright, E.; Neethirajan, S.; Warriner, K.; Retterer, S.; Srijanto, B. Single cell swimming dynamics of Listeria monocytogenes using a nanoporous microfluidic platform. Lab Chip 2014, 14, 938–946.

28. Sanders, L.; Andermann, T. M.; Ottemann, K. M. A supplemented soft agar chemotaxis assay demonstrates the Helicobacter pylori chemotactic response to zinc and nickel. Microbiology 2013, 159, 46–57.

29. Tan, S.; Berg, D. E. Motility of urease-deficient derivatives of Helicobacter pylori. J. Bacteriol. 2004, 186, 885–888.

30. Spohn, G.; Scarlato, V. Motility, Chemotaxis, and Flagella. In Helicobacter pylori: Physiology and Genetics·, Mobley, H. L., Mendz, G. L., Hazell, S. L., Eds.; Washington (DC): ASM Press, 2001; Chapter 21.

31. Nolan, K. J.; McGee, D. J.; Mitchell, H. M.; Kolesnikow, T.; Harro, J. M.; O'Rourke, J.; Wilson, J. E.; Danon, S. J.; Moss, N. D.; Mobley, H. L. T.; Lee, A. In vivo behavior of a Helicobacter pylori SS1 nixA mutant with reduced urease activity. Infect. limmun. 2002, 70, 685–691.

32. McGee, D. J.; Radcliff, F. J.; Mendz, G. L.; Ferrero, R. L.; Mobley, H. L. Helicobacter pylori rocF is required for arginase activity and acid protection in vitro but is not essential for colonization of mice or for urease activity. J. Bacteriol. 1999, 181, 7314–7322.

33. Huang, J. Y.; Sweeney, E. G.; Sigal, M.; Zhang, H. C.; Remington, S. J.; Cantrell, M. A.; Kuo, C.J.; Guillemin, K.; Amieva, M.R. Chemodetection and destruction of host urea allows Helicobacter pylori to locate the epithelium. Cell Host Microbe 2015, 18, 147–156.

